# Morphological evolution and phylogeny are unlinked in Hawaiian *Doryopteris* (Cheilanthoideae: Pteridaceae)

**DOI:** 10.64898/2026.10.06.757157

**Authors:** Priscilla Lau, Xuening Zhang, Jacob B. Landis, J. Gordon Burleigh, Layne Huiet, Kenneth R. Wood, Miles K. Thomas, Rosana Zenil-Ferguson, Carl J. Rothfels, Carrie M. Tribble

## Abstract

**Premise of the study:** *Doryopteris* J.Sm. is a genus of dry-adapted ferns found widely across the southern hemisphere and the tropics. In the Hawaiian Islands, the genus is represented by four endemic species and one putative hybrid. Two of the endemic species (*D. decora* and *D. decipiens*) are widespread across all major islands, while two (*D. angelica* and *D. takeuchii*) are federally and state-listed endangered single-island endemics.We aimed to resolve the evolutionary relationships and biogeographic history of Hawaiian *Doryopteris*.

**Methods:** W used a target-capture approach and the GoFlag 408 flagellate land plant probe set to sequence more than 400 nuclear loci for 40 Hawaiian and 29 non-Hawaiian individuals in *Doryopteris* and the related genera *Lytoneuron* and *Ormopteris*. After processing the sequences into alignments and single nucleotide polymorphism sites, we performed phylogenetic inference and population-level principal component analyses.

**Key results:** We infer that Hawaiian *Doryopteris* are monophyletic and are nested within the pantropical species *D. concolor*. Within Hawai‘i, clades and clusters of individuals correspond to geographical, geological, and climatological factors, more so than species identifications, with a few exceptions. We posit that Hawaiian *Doryopteris* represents an incipient radiation at the very earliest stages of divergence, and that budding speciation has led to non-monophyly of several taxa.

**Conclusions:** Our inference using genome-wide sequencing data sheds light on the complex evolutionary history of Hawaiian *Doryopteris*. Overall, the phylogeny of Hawaiian *Doryopteris* shows strong links with geographical, geological, and climatological processes and little with morphological evolution.

## Introduction

The evolutionary processes mediating how organisms disperse to and subsequently establish and diversify in new habitats determine how biodiversity patterns emerge and vary across space and time. The geographical isolation of islands enhance the effects of these processes, creating ideal systems for studying the formation of biodiversity (Whittaker et al. 2017, Warren et al. 2015, Carlquist 1974). Genomic tools have the potential to revolutionize the study of island-based radiations by revealing cryptic diversity and the underlying mechanisms of phenotypic variation and adaptation, yet few studies have applied such tools to island lineages (Cerca et al. 2023).

The Hawaiian archipelago has formed via movement of the Pacific Plate over a volcanic hotspot, which creates a chronosequence of islands. These islands range from submerged seastacks—the oldest is Meiji Seamount, which formed around 80 million years ago (mya) and is now almost 6,000 km away from the youngest island—to newly emerging volcanism above and below the ocean’s surface on Hawai‘i Island and the subsurface volcano Kama‘ehuakanaloa (Carson & Clague 1995). The modern “high islands” formed approximately 5.1 mya (Ni‘ihau, the furthest northwest) to less than 0.5 mya (Hawai‘i Island, the furthest southeast), and Hawai‘i Island contains several active volcanoes, including Kīlauea, one of the most active volcanoes on Earth (Carson & Clague 1995, Price & Clague 2002). The staggered formation of discrete volcanoes, the connection and disconnection of these volcanoes through shifting sea levels, and the erosion and sinking of land have shaped a matrix of (dis)connectivity, all in the context of extreme isolation from the mainland or even other islands: the Hawaiian islands are approximately 4,000 km away from any other high islands or continents, making them one of the most isolated places on Earth (Price 2004). The Hawaiian high islands, referred to as the Hawaiian Islands hereafter, form a tropical archipelago with varied microclimates, including desert, tropical alpine, rain forest, and dry forest zones (Edwards-Calma et al. 2024, Wagner et al. 1999). Within this climatological and geological context, several Hawaiian lineages have become textbook examples of plant adaptive radiations—bursts of rapid lineage diversification along with phenotypic adaptation—including the Hawaiian silversword alliance (Baldwin et al. 2026, Baldwin & Sanderson 1998, Witter & Carr 1988), the Hawaiian lobelioids (Rose et al. 2025, Givnish et al. 2004), and Hawaiian *Cyrtandra* (Kleinkopf et al. 2019, Cronk et al. 2005).

The Hawaiian flora comprises approximately 1,300 native vascular plant species, of which 90% are endemic (Wagner et al. 1999, Imada et al. 2025, Wagner et al. 2026). Some elements of this flora, such as those lineages mentioned above, have received considerable attention from botanists and evolutionary biologists over the past century. These lineages, all angiosperms, share in common high levels of extant diversity from the accumulation of new lineages and rapid phenotypic adaptations to novel niches. In contrast, we know relatively little about other elements of the Hawaiian flora. In particular, few studies have focused on non-flowering plants—though see Ranker et al., *e.g.*, 2016, 1994, 2019, and recent studies by Mendez-Reneau et al. (2024), among others. The Hawaiian fern flora—described in Ranker (2016) and Palmer & Thomas (2027)—is dominated not just by species-rich radiations, but also by a series of dispersal and establishment events with varied diversification outcomes. These diversification outcomes range from endemic radiations (*e.g.*, the 10 species of the endemic genus *Adenophorus*), endemism of a single species with closest relatives outside of Hawai‘i (*e.g.*, *Marsilea villosa*, ‘ihi‘ihi), and the apparent lack of speciation due to ongoing gene flow with non-Hawaiian populations (*e.g.*, the indigenous species *Asplenium adiantum-nigrum*, which has putatively established in Hawai‘i up to 17 times; Ranker et al. 1994).

Species complexes are relatively common in the Hawaiian flora (Palmer & Thomas 2027, Wagner et al. 1999). In these groups, morphological variation forms a continuous gradient, or gradates between murky clusters, rather than falling into discrete taxonomic units. When applying molecular techniques to such complexes, studies have demonstrated that ancestral polymorphism, extensive hybridization, and gene flow contribute to morphological variation across populations, despite limited molecular divergence that has occurred in the very recent past (Choi et al. 2021, Stacy et al. 2014, Mendez-Reneau et al. 2024). For example, *Metrosideros polymorpha* (‘ō hi‘a lehua), a dominant forest tree and keystone species, occupies an impressive breadth of climatic conditions such as mesic forests, recent lava flows, and moist forests across low to high elevations in the Hawaiian Islands (Wagner et al. 1999). Perhaps unsurprisingly then, the eight *M. polymorpha* varieties (Wagner et al. 2026, Imada et al. 2025) display remarkable phenotypic variation; Choi et al. (2021) note that the variation in leaf mass per area of Hawai‘i Island *M. polymorpha* is comparable to that of all woody plant species globally. This variation is thought to have arisen from ancestral polymorphism that facilitated rapid phenotypic adaptation despite porous reproductive barriers between taxa in an incipient radiation (Choi et al. 2021). Mendez-Reneau et al. (2024) tackled a separate lineage: *Polypodium pellucidum* and its zero to four infraspecific taxa, depending on taxonomic authority (Imada et al. 2025, Wagner et al. 2026, Palmer & Thomas 2027, Palmer 2003). They found evidence for extensive hybridization across the varieties, contributing to morphological complexity and taxonomic uncertainty.

Much like the above-described species complexes, Hawaiian members of the fern genus *Doryopteris* J.Sm. (Pteridaceae, Cheilanthoideae, ‘iwa‘iwa, kumuniu) display morphological variation that has confounded their classification into discrete species (Fig. 1, Palmer & Thomas 2027, Yesilyurt 2004). *Doryopteris* comprises approximately 25 taxa distributed across the tropics. Central and South America contain the highest taxon richness, but four to six taxa (depending on taxonomic authority) occur in Hawai‘i, and one species (*D. concolor*) occurs pantropically including in tropical Africa, Southeast Asia, the South Pacific, and South and Central America (Yesilyurt 2023). Two African taxa (*D. kirkii* and *D. nicklesii*) have been variably treated as part of this widespread *D. concolor sensu lato*, and we adopt this broad circumscription in this manuscript. *Doryopteris* species are typically differentiated by leaf characters, such as size, presence of wings between the two lowest pinna pairs, shape of ultimate segments, and distance between first and second pinna pairs (Palmer 2003). The Hawaiian taxa include four widely recognized species (*D. angelica* K.R.Wood & W.H.Wagner, *D. takeuchii* (W.H.Wagner) W.H.Wagner, *D. decora* Brack., and *D. decipiens* (Hook.) J.Sm.), one putative hybrid combination (*D. subdecpiens*, *D. decipiens × D. decora*), and one contested species (*D. tryonii* O.Deg. & I.Deg., recognized by Yesilyurt 2005, but not by Wagner et al. 2026, Imada et al. 2025, or Palmer & Thomas 2027). Two species, *D. decipiens* and *D. decora*, and their putative hybrid *D. subdecipiens*, are frequently described as widespread across the Hawaiian islands, while *D. angelica* and *D. takeuchii* are both narrow single-island endemics restricted to small populations on Kaua‘i and O‘ahu respectively (Fig. 1, Palmer & Thomas 2027).

**Figure 1:**
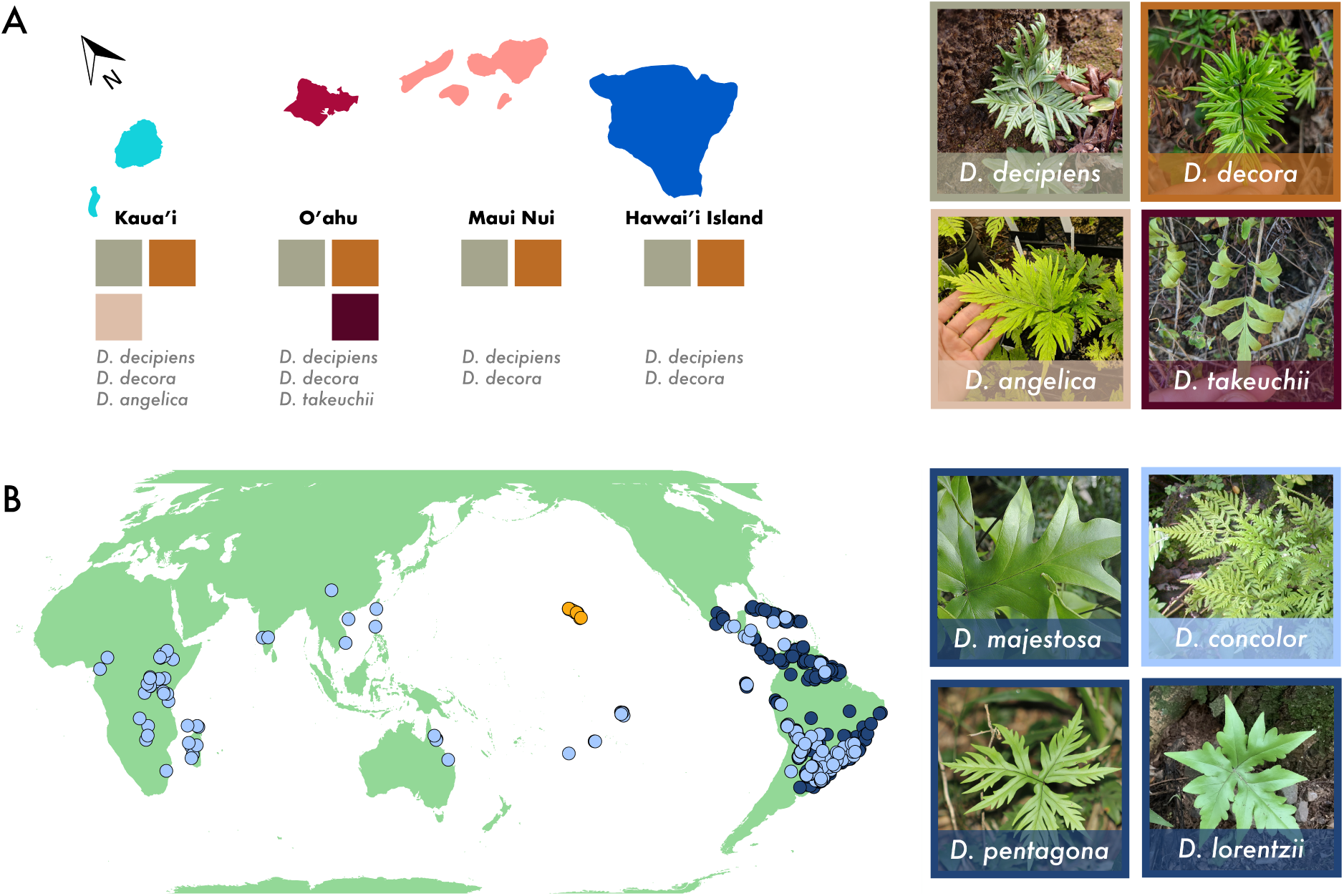
Distribution and morphological diversity of *Doryopteris*. (A) Hawaiian distribution and diversity of *Doryopteris*. Map of the main Hawaiian islands (color coded by island clusters) and the presence/absence of Hawaiian *Doryopteris* spp. on each main island group, and on the right photographs of the four recognized non-hybrid species in Hawai‘i. All photographs by CMT. (B) Global distribution and diversity of *Doryopteris*. Hawaiian species are indicated in yellow, *D. concolor s.l.* in light blue, and all other *Doryopteris* spp. in dark blue. To the right, photographs of select non-Hawaiian *Doryopteris* spp. (right). Photograph of *D. concolor* by KW, *D. lorentzii* by R. M. Guerra on iNaturalist, *D. pentagona* by Romi Galeota Lencina on iNaturalist, and *D. majestosa* by Dick Culbert from Flickr.

Yesilyurt (2004) revised the taxonomy of Hawaiian *Doryopteris* using morphological data, but this taxonomy was not fully accepted in subsequent treatments of the Hawaiian flora (Imada et al. 2025, Wagner et al. 2026, Palmer & Thomas 2027). Three molecular phylogenetic studies have included Hawaiian *Doryopteris* (Yesilyurt et al. 2015, Yesilyurt 2023, Schuettpelz et al. 2025). Each included a sample of *D. decora*, which was consistently supported as closely related to the widespread and pantropical *D. concolor*. Yesilyurt (2023) additionally included a sample of *D. decipiens*, which was resolved in *Lytoneuron* rather than with the sampled individual of *D. decora*; this result is likely due to labeling error or contamination (Yesilyurt 2023). The putative hybrid status of *D. subdecipiens* was determined based on its intermediate morphology between *D. decora* and *D. decipiens*, as these species form a spectrum of continuous variation that defies clear classification (Palmer 2003, Yesilyurt 2005). However, we have observed populations of *D. subdecipiens* without any nearby populations of *D. decipiens* or *D. decora*, for example on the island of Lāna‘i (unpublished), and specimens examined of *D. subdecipiens* do not contain aborted spores (Palmer & Thomas 2027). To date, no study has clarified the relationships among Hawaiian *Doryopteris*, confirmed whether Hawaiian *Doryopteris* represents the result of a single or multiple dispersal events to Hawai‘i, evaluated the hybrid status of *D. subdecipiens* with molecular data, or examined island-level relationships within the widespread species.

*Doryopteris*, like many members of the Cheilanthoideae subfamily, typically occupy dry to mesic habitats, and this is also true of the Hawaiian taxa (Tryon 1942, Yesilyurt et al. 2015, Edwards-Calma et al. 2024, Palmer & Thomas 2027). Their affinity for tropical dry ecosystems, one of the most threatened habitats worldwide (Stan et al. 2024, Miles et al. 2006), has led to conservation concerns within Hawai‘i. Both *D. angelica* and *D. takeuchii* are federally recognized under the Endangered Species Act, and NatureServe lists *D. decipiens* as G3, “Vulnerable” globally (Edwards-Calma et al. 2024, NatureServe 2026). *Doryopteris angelica*, restricted to Kaua‘i, is the only species of *Doryopteris* with an assessment by the International Union for the Conservation of Nature (IUCN) and was assessed as “Critically Endangered”, the only step above “Extinct in the Wild” (Edmonds & Walsh 2015). The taxonomic uncertainties described in the paragraph above further complicate attempts to assess and conserve *Doryopteris* in Hawai‘i, as without a stable and reliable taxonomy, it is not possible to determine species distributions or rarity.

Recent developments in next-generation sequencing technology have opened the door to generating genome-scale data from herbarium specimens, even when DNA from those specimens is fragmented and degraded (Funk 2018, Folk et al. 2021, Bieker & Martin 2018). Sequencing plant tissue from herbarium specimens avoids further sampling pressure on highly sensitive plant species or populations, such as *D. angelica* and *D. takeuchii*, making it particularly appealing for studies of threatened or endangered taxa. The GoFlag probe set targets over 400 single copy nuclear loci shared across flagellate plants, enabling targeted next-generation sequencing for lineages that lack genomic resources (Breinholt et al. 2021). Data from the GoFlag project have been used to successfully resolve phylogenies across deep and shallow nodes of the tree of life, even in the face of hybridization (*e.g.*, Fawcett et al. 2021, Mendez-Reneau et al. 2024, Tseng et al. 2024).

In this study, we employed the GoFlag probe set to infer the phylogenetic relationships within Hawaiian *Doryopteris* and across *Doryopteris* spp. in order to (1) provide a robust backbone phylogeny of *Doryopteris*, (2) determine if Hawaiian *Doryopteris* is the result of one or multiple dispersal events to the Hawaiian Islands, and (3) evaluate support for the current morphology-based classification of Hawaiian *Doryopteris*, including the putative hybrid status of *D. subdecipiens*.

## Materials and Methods

### Data collection

We sampled plant tissue from 69 individuals, including both silica-dried samples and herbarium specimens (see Supplemental Materials Table S1). This sample includes a dense representation of Hawaiian *Doryopteris*—40 individuals collected in Kaua‘i, Hawai‘i, Maui Nui, and O‘ahu—and 15 individuals of *D. concolor*, a widespread species that has been hypothesized to be closely related to Hawaiian *Doryopteris* (Yesilyurt et al. 2015). We also sampled seven non-*concolor Doryopteris* from outside of Hawai‘i, and seven samples of *Lytoneuron* and *Ormopteris* as an outgroup.

We extracted DNA using a standard CTAB protocol (Doyle 1991). The extracted DNA was sent to RAPiD Genomics (Gainesville, FL) for library preparation, target enrichment, and sequencing. We used the GoFlag probe set to target 408 exons (hereafter referred to as loci) from 229 single- or low-copy nuclear genes (Breinholt et al. 2021), and sequenced the samples with 150 base-pair paired-end reads. We inspected the Hawaiian individuals and revised species identifications from herbarium specimens and specimen images using Palmer (2003) and an early draft of Palmer & Thomas (2027). Given the differences in taxonomic treatments for Hawaiian *Doryopteris* and the challenges in differentiating *D. subdecipiens* from its putative parents (*D. decora* and *D. decipiens*), we took a conservative approach, labeling specimens as *D. subdecipiens* unless they very cleanly fell within the descriptions of *D. decora* or *D. decipiens*. If a specimen was not available, or if the individual did not fall within *Doryopteris* in resulting phylogenies, we removed it from all downstream analyses.

### Phylogenetic tree inference

We used an approximate multispecies coalescence approach to infer the phylogeny of *Doryopteris* from the alignments of the targeted loci (*i.e.*, excluding flanking regions). First, we assembled the raw reads into sequences using the GoFlag-specific pipeline described in Breinholt et al. (2021). Then, we aligned the sequences in each locus using MAFFT (Katoh et al. 2019) under default settings and trimmed the alignments using trimAl (Capella-Gutiérrez et al. 2009) under the “strictplus” condition. We inspected the alignments and manually adjusted any segments that were obviously shifted and non-homologous.

We inferred locus trees under the default setting (HKY model with empirical base frequencies) in IQ-TREE 2 (Minh et al. 2020) with 1000 rapid bootstrap replicates. In some cases, the GoFlag assembly pipeline outputs more than one sequence per sample for a locus, and so some locus trees contain multiple tips representing the same individual. Following the steps below, we resolved these instances such that each locus tree contained at most one copy per individual.

1. If the majority of individuals had multiple copies, and copies across individuals formed monophyletic clades (e.g., likely representing an ancestral locus duplication event), we split the locus into multiple alignments and re-inferred a phylogeny from each of the resulting loci.
2. If an individual contained multiple copies and the copies are monophyletic, we randomly pruned all but one copy from the tree. If the copies were not monophyletic, we excluded all copies of that individual from the locus.
3. If we could not prune out at least one monophyletic subtree with no more one copy per individual, we excluded the locus from subsequent analyses.

We filtered out loci with fewer than 12 individuals, resulting in 316 loci for the downstream analyses. We realigned the loci using MAFFT and re-inferred the locus trees using IQ-TREE 2. We collapsed the nodes with lower than 10% bootstrap support in each locus tree. From the locus trees, we inferred the species phylogeny using ASTRAL-II (Mirarab & Warnow 2015), where each individual was treated as its own “species”.

To provide more resolution within the Hawaiian samples, we also zoomed in on a subset of samples. We determined the clade of closest relatives to the Hawaiian samples from the phylogeny inferred above. We then retained only those “outgroup” samples and the Hawaiian samples, applying two sets of filters. In our first approach, we excluded all individuals of *D. subdecipiens*, as hybrids can cause topological instability (Schneider et al. 2016, McDade 1992), and we excluded all individuals represented in fewer than 50% of the gene trees. This pruning step resulted in excluding 12 *D. subdecipiens* samples and three additional Hawaiian samples (two *D. decipiens* and one *D. takeuchii*). In our second approach, we tested the impact of retaining *D. subdecipiens* and only excluded samples represented in fewer than 50% of gene trees (the two *D. decipiens* and one *D. takeuchii*). For each approach, we re-estimated alignments with the reduced set of samples, estimated gene trees as described above, and inferred a species tree. Unlike the species-tree methods described above, here we retained poorly supported nodes in the gene trees and inferred the species tree using weighted ASTRAL (wASTRAL) implemented in the ASTER package (Zhang et al. 2025, Zhang & Mirarab 2022).

### SNP calling and processing

Characterized by low branch support and short branch lengths, our inferred phylogenies did not show clear structure within the Hawaiian clade. To explore these results further, we performed some population-level analyses based on single-nucleotide polymorphism (SNP) data. We included all Hawaiian individuals and the closely related Pacific *D. concolor* clade (Clade B) as an outgroup.

First, we processed raw reads of both targeted and flanking regions using fastp v0.20.0 (Chen et al. 2018). This procedure used auto-detection to remove adapters, trim polyG strings, and filter out reads shorter than 60 bp or with a quality score lower than 20. Then, we mapped processed reads to one of two reference files using bwa-mem2 v2.0pre2 (Vasimuddin et al. 2019): the probe sequences or the full locus (including flanking regions), both generated from the sequence output of the GoFlag pipeline randomly pruned to retain one sequence per individual per locus. We converted the output SAM files to BAM files and sorted the output using samtools v1.16.1 (Danecek et al. 2021). After marking duplicates, we called SNPs using HaplotypeCaller, GenomicsDBImport, and GenotypeGVCFs in GATK v4.3.0 (Poplin et al. 2017, Van der Auwera & O’Connor 2020). Specifically, we performed two separate joint genotyping steps—one including the variants only, and one including non-variants given the differences in branch lengths observed in previous studies (Suissa et al. 2024).

We filtered variants using VCFtools v0.1.16 (Danecek et al. 2011), keeping only biallelic sites with (i) not more than 40% missing data across samples, (ii) depth between 10 and 100, and (iii) a minor allele count of at least three. We removed samples with more than 50 percent missing data as calculated by VCFtools to obtain a set of high quality SNPs. This resulted in a SNP dataset of 25 Hawaiian individuals and five *D. concolor* individuals. We note that unfortunately the sole individual from Hawai‘i Island was filtered out in this step. Finally, we filtered out sites in linkage disequilibrium (LD) using PopLDdecay v3.40 (Zhang et al. 2019) and plink2 v3.7 (Chen et al. 2019) with a r2 value of 0.2 in a 10 kb window with a 5 kb step size. This filtering resulted in 47943 SNPs for the dataset that excluded the outgroup and 48645 SNPs for the dataset that included the outgroup before LD pruning, and 4466 and 5276 SNPs for the two datasets after LD pruning. For downstream analyses, we converted filtered VCF files to fasta, nexus, and binary nexus using vcf2phylip v2.8 (Ortiz 2019).

### Population-level analyses

To visualize the variation in genomic composition across individuals, we performed principal component analyses (PCA) on the SNP data, both including and excluding non-Hawaiian individuals, using the R package SNPrelate (Zheng et al. 2012). In addition, we inferred unrooted phylogenetic pseudonetworks using the neighbor-net method in SplitsTree v4 (Huson & Bryant 2006).

## Results

### Species identifications

After examining specimens for all Hawaiian individuals, we changed the names of 13 samples (Tab. 1). We identified five specimens originally determined as *D. decipiens* as *D. subdecipiens*, four specimens originally determined as *D. decora* as *D. subdecipiens*, and two specimens originally determined as *D. subdecipiens* as *D. decora*. We identified two specimens only determined to genus (*Doryopteris sp.*) as *D. decipiens* and *D. angelica*. All other specimens retained their original identifications. Notably, these reidentifications resulted in only one *D. decipiens* sample from Kaua‘i and no sampled *D. decora* from Maui Nui or O‘ahu, raising questions regarding the supposed Hawai‘i-wide distributions of these two species, though thorough examination of herbarium specimens beyond those included in our study would be required to confirm this suspicion.

**Table 1:** Taxonomic reclassification of specimens. All specimens not included in this table retained their original identification.

| Original ID | New ID | Locality | Collection Number |
| --- | --- | --- | --- |
| *D. decipiens* | *D. subdecipiens* | Hawai'i Island | C Morden 1119 |
| *D. decipiens* | *D. subdecipiens* | Oahu | DD Palmer 763 |
| *D. decipiens* | *D. subdecipiens* | Kahoolawe | KR Wood 12923 |
| *D. decipiens* | *D. subdecipiens* | Oahu | KR Wood 10713 |
| *D. decipiens* | *D. subdecipiens* | Kauai | KR Wood 18265 |
| *D. decora* | *D. subdecipiens* | Oahu | DD Palmer 764 |
| *D. decora* | *D. subdecipiens* | Oahu | KR Wood 10715 |
| *D. decora* | *D. subdecipiens* | Kauai | DD Palmer 584 |
| *D. decora* | *D. subdecipiens* | Kauai | KR Wood 18028 |
| *D. sp.* | *D. decipiens* | Maui | C Morden 2138 |
| *D. sp.* | *D. angelica* | Kauai | DD Palmer 2178 |
| *D. subdecipiens* | *D. decora* | Kauai | KR Wood 17847 |
| *D. subdecipiens* | *D. decora* | Kauai | KR Wood 15883 |

### Phylogenetic tree inference

We find strong support for the backbone topology of *Doryopteris*. Specifically, *D. concolor* and Hawaiian individuals form a clade sister to the rest of *Doryopteris* (Fig. 2), and *Doryopteris* is sister to *Lytoneuron* + *Ormopteris*. Within clade C, “other *Doryopteris*”, we infer the sister species *D. pentagona* and *D. lorentzii* as sister to the rest of clade C. We infer sister relationships between (i) *D. nobilis* and *D. patula*, and (ii) *D. varians*, including an individual identified as *D.* cf. *varians*, and *D. collina*. These results are all maximally supported (posterior probability, *pp*, of 1.0).

**Figure 2:**
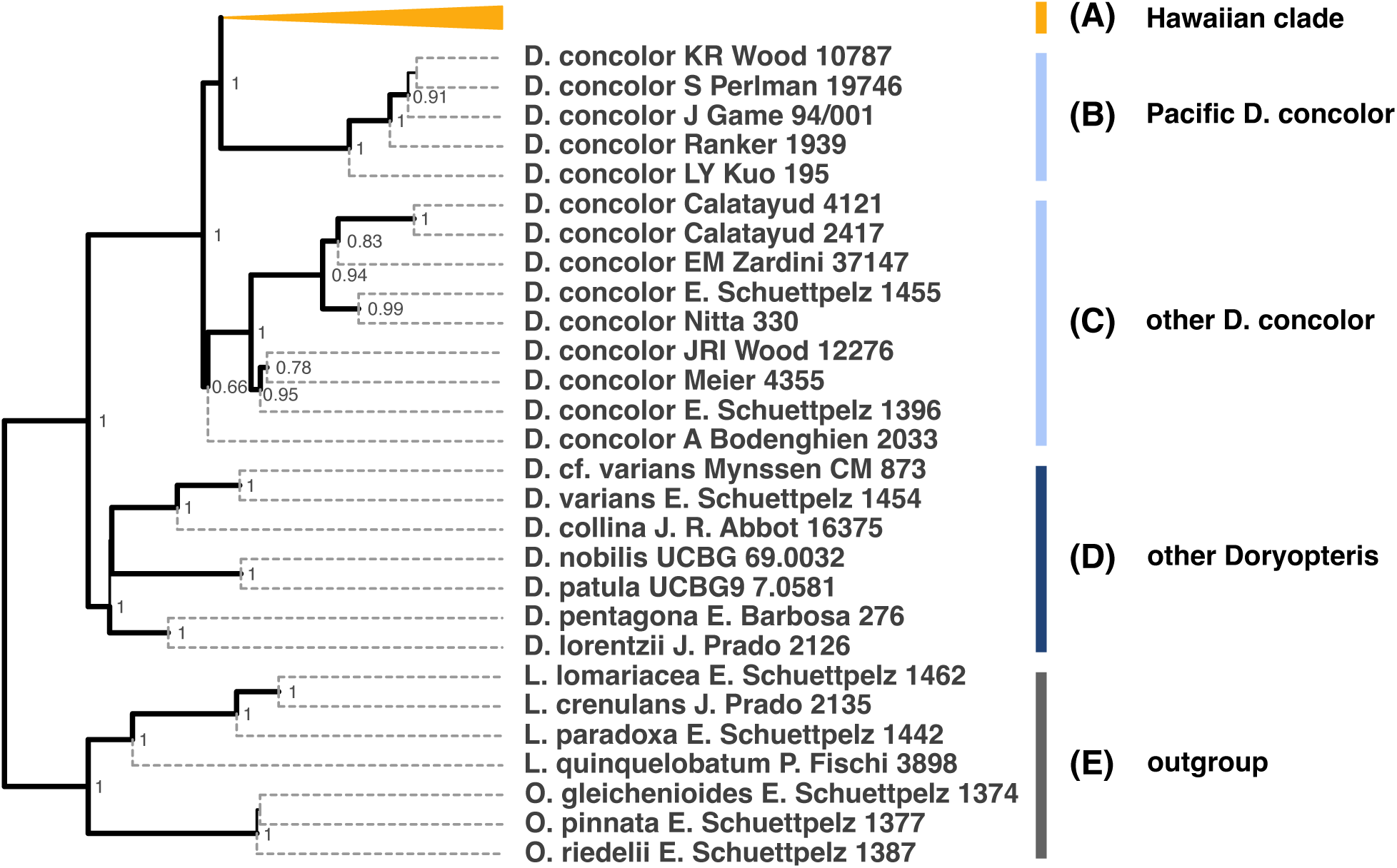
Backbone relationships among *Doryopteris* inferred using 316 exon regions from the GoFlag probe set with ASTRAL. Dashed gray lines represent terminal branch lengths, which are not shown because their lengths are not estimated by ASTRAL. The Hawaiian clade is collapsed. Branch thickness (excluding tip branches) represents posterior support. Posterior support *>* 0.6 is displayed. Major non-Hawaiian clades are labeled following the color code in Fig. 1

Within the *concolor*–Hawaiian clade, we recover Hawaiian *Doryopteris* as monophyletic and nested within *D. concolor* (Fig. 2). The Hawaiian individuals are most closely related to Clade B: *D. concolor* from south Pacific islands (Cook Islands, the Marquesas Islands, and French Polynesia) and the northwest Pacific (Taiwan). Clades A + B (Hawaiian *Doryopteris* and Pacific *D. concolor*) are collectively sister to a clade of continental *D. concolor* primarily from South America, clade C. One South Pacific sample (Nitta 330, from Mo‘orea), is nested within clade C. Our single sample from tropical Africa *(D. concolor* Bodenghien 2033) is sister to the South American clade, albeit on a very short branch with poor support (*pp* = 0.66).

We obtain relatively high posterior support for the splits among *D. concolor* individuals, as well as the split between Hawaiian individuals and Pacific *D. concolor* (*pp >* 0.6); however, the splits within the Hawaiian clade generally have low posterior support (Fig. 3). Furthermore, we do not observe a clear association between the morphology-based taxonomic assignment of the individual and its phylogenetic position within the Hawaiian clade, particularly with regards to *D. subdecipiens*, which is scattered across the tree. However, when we exclude *D. subdecipiens* (Fig. 3, left), we are able to identify general patterns. A poorly-supported node (*pp <* 0.6) splits the Hawaiian clade into two geographically defined subclades: Kaua‘i vs. O‘ahu + Maui Nui. Within the Kaua‘i clade, *D. angelica* is (almost) monophyletic: a single *D. decipiens* individual (KR Wood 17843) is nested within it. Similarly, that *angelica* clade is (almost) sister to *D. decora*; a single isolated *decora* sample (DD Palmer 585) renders the second species non-monophyletic. Within the O‘ahu and Maui Nui clade, our single representative of *D. takeuchii* is nested within *D. decipiens*. When we add in the *D. subdecipiens* individuals, these clades largely remain but with *D. subdecipiens* individuals interspersed with Kaua‘i *D. decora* and with O‘ahu and Maui Nui *D. decipiens* (Fig. 3, right). The only sample that switches clades when *D. subdecipiens* is included is DD Palmer 585 (*D. decora* from Kaua‘i), which is sister to the Kaua‘i clade when *D. subdecipiens* is excluded but sister to the O‘ahu and Maui Nui clade when *D. subdecipiens* is included, rendering Kaua‘i non-monophyletic. In both trees, the placement of DD Palmer 585 is poorly supported (pp *<* 0.6).

**Figure 3:**
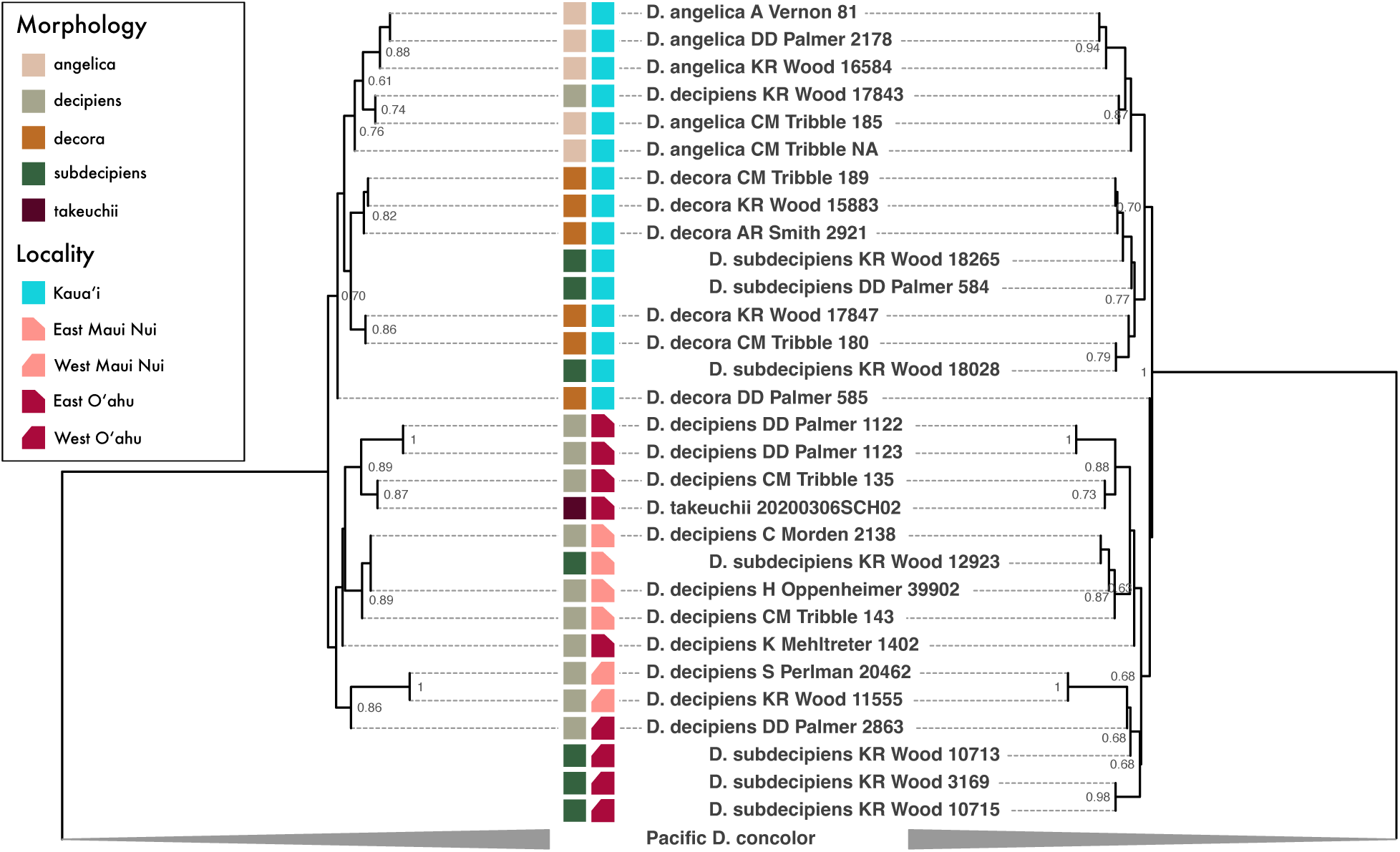
Phylogenetic trees of Hawaiian *Doryopteris* using 302 (left) and 305 (right) aligned target regions from the GoFlag probe set, inferred with wASTRAL. Dashed lines represent terminal branch lengths, which are not shown because their lengths are not estimated by wASTRAL. (Left) The phylogeny estimated with *D. subdecipiens* samples excluded; (Right) The phylogeny estimated with *D. subdecipiens* samples included. Posterior support *>* 0.6 is displayed. Within the Hawaiian clade, colored boxes show the morphological identifications and island locality of each sample, respectively.

### Population-level analyses

Our population-level analyses that include outgroup samples show that the Hawaiian *Doryopteris* are genetically distinct from *D. concolor* in the south and northwest Pacific (Fig. 4B,D,F). In the PCA, most of the variation between Hawaiian and non-Hawaiian individuals can be explained by the first eigenvector (25.63%), while the second eigenvector (6.11%) mostly explains the variation among Hawaiian individuals. We find a similar pattern in the distance-based phylogenetic network, where Hawaiian and non-Hawaiian individuals are separated by long branches (Fig. 4).

**Figure 4:**
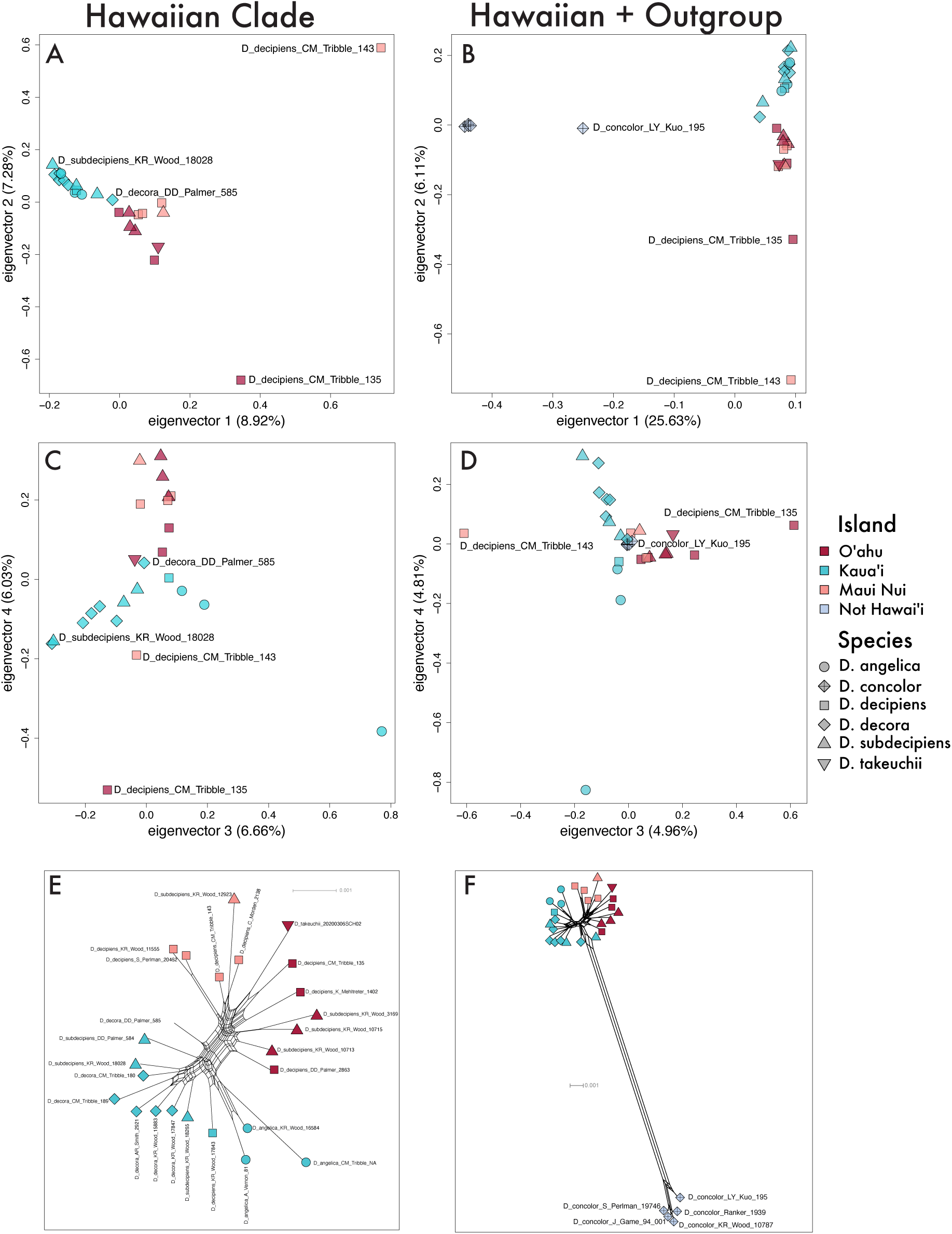
Population-level patterns of the Hawaiian clade and outgroup samples (*D. concolor* from the Pacific) using SNPs called from flanking and target regions. Panels A–D show principal component analyses of the Hawaiian clade (A, C) and Hawaiian clade + outgroup (B,D). Panels E (Hawai‘i only) and F (Hawai‘i and outgroup) show a splitstree pseudonetwork visualization of the same two datasets. Individuals are colored by geography and shapes correspond to species.

Our analyses that exclude non-Hawaiian individuals provide higher resolution of the genetic variation within Hawaiian *Doryopteris* (Fig. 4A,C,E). In the PC analysis, individuals are generally clustered by their island locality (indicated by color) on the first two eigenvectors. The third and fourth eigenvectors show some evidence of clustering by species (indicated by shape). Within Kaua‘i, *D. angelica* is differentiated from *D. decora* and *D. subdecipiens* along eigenvector 3. In contrast, O‘ahu and Maui Nui individuals are interspersed along eigenvector 4, which appears to distinguish *D. decipiens* (negative values) from *D. subdecipiens* (positive values). However, the first two eigenvectors explain only 15.2% of the variation, indicating a lack of clear linear directionality in the data. In the phylogenetic network, we recover two main clusters: one contains exclusively individuals from Kaua‘i, while the other contains individuals from O‘ahu and Maui Nui. There are some subclusters of O‘ahu and Maui Nui samples, but these are not as distinct.

## Discussion

We set out to resolve a backbone phylogeny of *Doryopteris* using genome-scale nuclear data, to test the monophyly of Hawaiian *Doryopteris*, and to assess the relationships among Hawaiian *Doryopteris* taxa and within Hawaiian *Doryopteris* spp. across island populations. Our results represent the first genome-scale backbone phylogeny of *Doryopteris*. Our phylogeny is also the first peer-reviewed study to include representatives of multiple Hawaiian *Doryopteris* species, distribution-wide sampling of *Doryopteris concolor*, or multiple individuals of the four recognized Hawaiian *Doryopteris* spp. and putative hybrid *D. subdecipiens*. Our inferences clearly demonstrate that Hawaiian *Doryopteris* is the result of a single dispersal and establishment event, despite previous work suggesting the possibility of multiple dispersal events (Yesilyurt 2005, 2023). We also present a complex view of relationships among Hawaiian *Doryopteris*, one in which some individuals appear more closely related to individuals on the same island than their putative conspecifics on other islands. Below, we discuss these findings and their implications.

### The phylogeny of *Doryopteris* and related genera

We recover well-resolved backbone relationships within *Doryopteris*, largely concordant with a previous phylogenetic study based on two chloroplast markers (rbcL and trnLF; Yesilyurt et al. 2015). For example, in agreement with Yesilyurt et al. (2015), we infer Hawaiian *Doryopteris* to be closely allied with *D. concolor*, collectively forming a sister group to the rest of non-Hawaiian *Doryopteris*. Both studies also infer close relationships between *D. pentagona* and *D. lorentzii*, between *D. varians* and *D. collina*, and between *D. nobilis* and *D. patula* (called *D. majestosa* in Yesilyurt et al. 2015). However, we infer *D. pentagona* and *D. lorentzii* to be sister to the rest of clade C (other *Doryopteris*), contrary to Yesilyurt et al. (2015), which infers *D. lorentzii* and *D. pentagona* as collectively sister to a clade including *D. varians* and *D. collina*. In our tree, the branch subtending the split between *D. pentagona* + *D. lorentzii* from the remaining members of clade C is quite short, indicating rapid divergence between the respectively lineages. Our study leverages hundreds of nuclear markers compared to two chloroplast markers in Yesilyurt et al. (2015), so it is possible that either the scale of the data changes inferred relationships or that the nuclear vs. chloroplast genomes differ in their evolutionary histories. Unfortunately, our matK phylogeny (Supplemental Fig. S1) does not include sufficient sampling in clade C to allow us to differentiate between these two hypotheses. It is also possible that the position of the *D. lorentzii* and *D. pentagona* clade is sensitive to taxon sampling, as the Yesilyurt et al. (2015) phylogeny included roughly twice as many non-Hawaiian species as our present study.

### Distributions of Hawaiian *Doryopteris* spp

Our re-identification of Hawaiian *Doryopteris* resulted in a conspicuous absence of *D. decora* on O‘ahu and Maui Nui, and a paucity of *D. decipiens* on Kaua‘i (the only Kaua‘i *D. decipiens* is nested within *D. angelica*, discussed below). We took a conservative approach to identifying *D. decora* and *D. decipiens*, which appears to have limited the occurrences of these species from their previously understood, widespread distributions (as presented in Fig. 1). Our molecular results support the distinction of *D. decora* on Kaua‘i from individuals previously identified as *D. decora* on other islands, as the deepest split in our phylogeny separates Kaua‘i individuals from everything else. These results suggest a significant shift in our understanding of Hawaiian *Doryopteris* spp. and their distributions across the islands; perhaps *D. decora* and *D. decipiens* are not as widespread as previously thought, and are instead narrowly distributed, morphologically unique lineages found on one or a few islands (Kaua‘i and O‘ahu + Maui Nui respectively). Yesilyurt’s 2005 morphological treatment of *Doryopteris* in Hawai‘i provides some limited support in favor of this view. That study suggested that *D. tryonii* (a species not recognized in most current treatments of the Hawaiian flora, which instead treat it under *D. decora*) is restricted to Kaua‘i. A complete revision of Hawaiian *Doryopteris* is necessary in the future, as our results indicate that a detailed revision of these taxa is warranted.

### Paraphyletic *D. concolor* and the origin of Hawaiian *Doryopteris*

Hawaiian *Doryopteris* is nested within *D. concolor*, making *D. concolor* paraphyletic. Specifically, the *D. concolor* individuals we sampled are divided among two clades, one of individuals from south and northwest Pacific islands (clade B) and one composed primarily of samples from mainland South America (clade C). Clade C includes two samples from outside of South America: Nitta 330 from Mo‘orea and Bodenghien 2033 from the Democratic Republic of the Congo (discussed below). Our other individual from Mo‘orea (Ranker 1939) falls out in Clade B with other Pacific island individuals, so our result for Nitta 330 requires independent confirmation. The Hawaiian *Doryopteris* species are sister to clade B, and together this Pacific lineage is sister to the remaining *D. concolor* sampled in our analysis. *Doryopteris concolor* is an incredibly widespread species (see Fig. 1), and unfortunately our sampling does not include significant representation outside of the Americas and the Pacific. A single sample from the Democratic Republic of the Congo, Bodenghien 2033, is tentatively inferred within the South American *D. concolor* clade, but this relationship is poorly supported. Global sampling of *D. concolor*, including more sampling in tropical Africa, Southeast Asia, and Australia, and explicit sampling of the closely allied taxa (here treated as part of *D. concolor s.l.*: *D. kirkii* and *D. nicklesii*), would provide a more complete picture of the species’ spread across the tropics, and/or demonstrate the existence of cryptic species in this complex.

Based on our present sampling, it seems likely that *Doryopteris* originated in South America. Then, *D. concolor* branched out from the rest of *Doryopteris* and dispersed across the tropics, including the South Pacific islands. We infer the Pacific individuals to be phylogenetically close to the Hawaiian individuals, hinting at a northward dispersal from South Pacific islands to Hawaiian islands. However, the Taiwan sample of *D. concolor* (Kuo 195) is closer to the Hawaiian clade in our PC analysis (Fig. 4B), suggesting that more work is needed within *D. concolor* to resolve the dispersal history of this species and the potential dispersal route to the Hawaiian Islands from within the Pacific. Furthermore, we infer that Hawaiian *Doryopteris* is monophyletic, indicating that Hawaiian species are the result of a single dispersal event, in contrast to previous results suggesting otherwise (Yesilyurt 2005, 2023). Given the short branch lengths, unresolved relationships among Hawaiian individuals, and the young geological age of the current Hawaiian high islands (5.1 mya), the arrival of *Doryopteris* on Hawaiian islands is likely a recent event. This timing should be investigated in future studies.

### Dispersal and diversification across the Hawaiian Islands

We demonstrate that most of the current taxonomic definitions are not consistent with the evolutionary history of Hawaiian *Doryopteris*. Rather, our results suggest an ongoing, incipient radiation, in which morphologically distinct clusters are budding off from a widespread, generalized lineage within restricted geographical ranges. Below, we discuss the monophlyly (or lack thereof) of each Hawaiian *Doryopteris* species and evaluate the possibility of Hawaiian *Doryopteris* experiencing budding speciation and/or representing an incipient radiation.

*Doryopteris subdecipiens*, which represents intermediate morphologies between *D. decora* and *D. decipiens*, is non-monophyletic. We find little phylogenetic support for distinction between *D. subdecipiens* and *D. decora* within Kaua‘i, and between *D. subdecipiens* and *D. decipiens* within O‘ahu and Maui Nui, though this latter distinction is tentatively supported by the fourth eigenvector of our PC analysis (Fig. 4C). We do find consistent support for the differentiation of *D. angelica* from other *Doryopteris*, with the notable exception of a *D. decipiens* individual (Wood 17843), which was noted at the time of collection to resemble *D. angelica* in size and frond dissection, but which lacks *D. angelica*’s distinctive “winged” stipe; this specimen presumably represents a morphologically aberrant *D. angelica* individual. *D. angelica*’s climatic niche is generally more mesic than that of other members of Hawaiian *Doryopteris* (Edwards-Calma et al. 2024), further supporting its distinction from other lineages. This species is recovered either as nested within other Kaua‘i *Doryopteris* (Fig. 3left), thus suggestive of budding speciation, or as sister to a clade of Kaua‘i *D. decora* and *D. subdecipiens* (Fig. 3right), thus more suggestive of traditional, bifurcating speciation.

In contrast to the dissociation between current taxonomy (which largely reflects leaf morphology) and evolutionary relationships, we find substantial support that links the phylogenetic history to geology, geography, and climate, likely reflecting a history of dispersal events leading to incipient budding speciation across the Hawaiian Islands. We consistently recover a sister relationship between Kaua‘i and O‘ahu–Maui Nui individuals, with the possible exception of one Kaua‘i sample (*D. decora* Palmer 585) which switches position when *D. subdecipiens* is included. This sample is centrally located between island clusters in our PC analysis vectors 1 and 2 (Fig. 4A), corroborating its unstable position in the species tree. Therefore, we are unable to infer a likely ancestral island range for the initial establishment of *Doryopteris* in Hawai‘i. If Kaua‘i *Doryopteris* is truly paraphyletic as suggested in Fig. 3 (left), then the ancestral Hawaiian *Doryopteris* population could have found its root in Kaua‘i before dispersing southeast, a pattern consistent with the progression rule in Hawai‘i, in which species are hypothesized to have first arrived to the oldest high island (Kaua‘i), subsequently dispersing down the chain as more islands form (Funk & Wagner 1995).

We infer two clades of Maui Nui individuals nested within samples from O‘ahu. These two clades correspond to samples from the eastern and western sides of Maui island, consisting of two separate volcanoes: Mauna Kahālāwai (West Maui) and Haleakalā in the east. The one exception is a sample from Kaho‘olawe (KR Wood 12923), which is nested within the East Maui Nui clade. Kaho‘olawe, formerly part of Maui Nui but now a small island, is on the eastern side of the Maui Nui island group. These East and West Maui Nui clades are nested within O‘ahu in our species-tree analysis, suggesting multiple dispersal from O‘ahu down the chain (Fig. 3). The two Maui Nui clades are not only nested within O‘ahu, but are specifically each nested within subclades of O‘ahu that correspond to the eastern and western sides of that island, respectively (*i.e.*, the East Maui Nui clade is nested within the East O‘ahu clade, and the West Maui Nui clade is nested within the West O‘ahu clade). This result is somewhat surprising, as both East and West O‘ahu are physically closer to West Maui Nui than either is to East Maui Nui. However, the western sides of O‘ahu and Maui are both generally drier than the eastern sides due to the rain shadow from the prevailing trade winds, which come out of the northeast (Giambelluca 1986). It is tempting to suggest that affinity within eastern vs. within western populations of *Doryopteris* across these two islands reflects climatic preferences, but more focused niche modeling would be needed to formally test this hypothesis. (Edwards-Calma et al. (2024) did model the climate niches for Hawaiian *Doryopteris*, but in the absence of molecular data to suggest otherwise, that study combined samples of the same “species” regardless of locality).

Four of the five East O‘ahu Clade samples come from tuff cones formed during posterosional, second stage eruptive activity known as the Honolulu Volcanic Series, including Lē‘ahi (Diamond Head, one *D. takeuchii*) and Kohelepelepe (Koko Crater, three *D. decipiens*). The fifth sample (*D. decipiens*, Mehltreter 1402) is geographically close to the Kohelepelepe samples (from Kuli‘ou‘ou in the Ko‘olau range), but is more closely related to the East Maui Nui samples than to its nearest neighbors. The tuff cones of the Honolulu Volcanic series differ from the Ko‘olau range in three main ways. Tuff forms when volcanic ash cements as it cools, making it chemically and physically distinct from the basalt typical of most Hawaiian volcanoes (nepheline basanite and basanitoid magma vs. tholeiitic and olivine basalts; Macdonald et al. 1983). The tuff cones of the Honolulu Volcanic Series formed substantially later than O‘ahu’s primary volcanism period (approximately 1.3 mya difference in age) and have thus weathered less (Ozawa et al. 2005). Lastly, both Lē‘ahi and Kohelepelepe are generally drier parts of O‘ahu than even the leeward slopes of the Ko‘olau mountains, and are closer to coast. Thus, the relationships in our species tree suggest that clades within Maui Nui and especially O‘ahu could be budding off in response to fine-scale geological and climatological factors. It might come as a surprise to see such fine-scale differentiation of a spore-bearing lineage within a single island, as fern spores are thought to disperse widely on the wind (Tryon 1990). The maintenance of reproductive barriers in the face of such indiscriminate dispersal should be the subject of future research.

Regardless of the nature of these relationships, an affinity between O‘ahu and Maui Nui *Doryopteris* populations is perhaps unsurprising considering that these areas once formed one large continuous island, connected via a land bridge through much of what is now Moloka‘i (Carson & Clague 1995). In contrast, Kaua‘i has maintained an oceanic channel that separates it from the islands to the southeast since their formation (Carson & Clague 1995). The lack of samples from Hawai‘i Island is unfortunate, since Hawai‘i Island has also always maintained a sea channel separation between its nearest neighbor (Maui).

It is also possible that frequent introgression has swamped genetic signal of the distinction between species. For example, perhaps *D. subdecipiens* does represent hybrids between *D. decora* and *D. decipiens*. Under this explanation, we might expect that such hybrid individuals would fall out between the putative progenitors, in other words between the Kaua‘i and O‘ahu + Maui Nui clades. Instead, these individuals are well-supported in their respective island-clades (pp = 1.00, Fig. 3, right). Across all analyses, this geographic signal dominates, including in our wASTRAL phylogeny (Fig. 3), our SNP-based PCA (Fig. 4A,B), and our pseudonetworks (Fig. 4E,F). If introgression within islands has swamped out species-specific genetic signals, such introgression appears rampant enough that species boundaries must be quite porous.

### Conclusions and conservation implications

Our results paint a complex picture of *Doryopteris* evolution within Hawai‘i. Instead of four species and a hybrid, we suggest that Hawaiian *Doryopteris* represents a series of geographically, geologically, and/or climatologically unique populations or taxa that are in the process of budding off from a morphologically intermediate ancestral taxon (*D. subdecipiens*). Hawaiian *Doryopteris* represents an incipient radiation, a lineage in the very early stages of differentiation. These budding populations may result in fully distinct, sorted species, but our results suggest that this is not currently the case. Rather, we suggest a lava lamp model of speciation, one in which small blobs bud off from a large primary blob, float along more or less distinctively, before either rejoining the large blob or diverging fully. We are viewing a static picture of this lava lamp model. Alternatively, it is possible that rampant introgression has swamped out species-specific signal in our molecular data; such a conclusion implies that species boundaries are not being maintained in the face of gene flow and that geographic signal dominates over remnant genetic signal of phylogenetic history. Regardless of the eventual fate of these lineages, we must make taxonomic and management decisions now. Both *D. angelica* and *D. takeuchii* are federally listed endangered species. Our results support the continued recognition of *D. angelica*, though more focused work within Kaua‘i could resolve whether *D. decipiens* is present on the island. Unfortunately, our limited sampling of *D. takeuchii* prevents us from evaluating that species. Our results indicate that a revision of *D. decipiens*, *D. decora*, and *D. subdecipiens* is warranted. In particular, morphological differences may exist between *D. subdecipiens* on O‘ahu and Maui Nui vs. on Kaua‘i. Given the gradation of morphology across these three taxa, it is possible that infraspecific taxonomic status is more appropriate in a future taxonomic revision.

## Author Contributions

CMT and CJR conceived of the project. XZ, PL, JBL, and LGB generated and analyzed the data. PL, CMT, MKT, and CJR interpreted results. PL and CMT drafted the manuscript, and all authors provided feedback and edits to the final manuscript.

## Data and code availability

Raw reads will be made available on NCBI SRA upon acceptance of this manuscript. Intermediary data files and results are available on Zenodo at DOI: 10.5281/zenodo.23018707. All code is available on GitHub at https://github.com/prilau/hawaiian_doryopteris.

## Acknowledgments

This material is based upon work supported by the NSF Postdoctoral Research Fellowships in Biology Program under Grant No. 2109835 to CMT. Any opinions, findings, and conclusions or recommendations expressed in this material are those of the authors and do not necessarily reflect the views of the National Science Foundation. Funding for this project also came from the University of California, Berkeley, Department of Integrative Biology and NSF-DEB 2144011 awarded to CJR. The Ferns* Lab Database, Eric Schuettpelz, the University of California and Jepson Herbaria, the Joseph F. Rock Herbarium at the University of Hawai‘i at Mānoa, the Herbarium Pacificum at the Bishop Museum, and the Herbarium of the National Tropical Botanical Garden provided samples for this project. All T&E specimens were collected under T&E permits 260202100252-PLANT and DOFAW Endorsement No: I2542, and State of Hawai‘i Forest Reserve permit No. KPI-2026-424. We thank the State of Hawai‘i Department of Land and Natural Resources Division of Forestry and Wildlife for their support of this work. In particular, we thank Susan Ching for assistance in collecting *D. takeuchii*, Adam Williams for his assistance with *D. angelica* permitting, and Matthew Keir for his overall support of this project. We additionally acknowledge R. M. Guerra on iNaturalist, Romi Galeota Lencina on iNaturalist, and Dick Culbert on Flickr for photographs presented in Fig. 1. We thank Jefferson Prado for his assistance with identifications of *D. varians*.

