## Supplemental Materials for "Morphological evolution and phylogeny are unlinked in Hawaiian *Doryopteris* (Cheilanthoideae: Pteridaceae)"

### Supplemental Material

#### S1 Specimen table

**Table S1:** Specimen collection data for *Doryopteris*, *Lytoneuron*, and *Ormopteris* accessions.

| Genus | Species | Collection Number | Institution | Accession Number | Locality | Material |
|---|---|---|---|---|---|---|
| Doryopteris | decora | DD Palmer 585 | HAW | HAW41381 | Kaua'i, Hawai'i | herb |
| Doryopteris | decora | CM Tribble 180 | UC | NA- not yet accessioned | Kaua'i, Hawai'i | silica |
| Doryopteris | decora | CM Tribble 189 | UC | NA- not yet accessioned | Kaua'i, Hawai'i | silica |
| Doryopteris | decora | Flynn sn | NA | MO4856634 | Kaua'i, Hawai'i | herb |
| Doryopteris | decora | AR Smith 2921 | UC | UC1877815 | Kaua'i, Hawai'i | herb |
| Doryopteris | decora | KR Wood 17847 | PTBG | PTBG1000068644 | Kaua'i, Hawai'i | silica |
| Doryopteris | decora | KR Wood 15883 | PTBG | PTBG1000084101 | Kaua'i, Hawai'i | silica |
| Doryopteris | decipiens | KR Wood 17843 | PTBG | PTBG1000069070 | Kaua'i, Hawai'i | silica |
| Doryopteris | decipiens | KR Wood 9071 | UC | UC2026830 | Kaua'i, Hawai'i | herb |
| Doryopteris | decipiens | H Oppenheimer 39902 | HAW | HAW41393 | Maui, Hawai'i | herb |
| Doryopteris | decipiens | S Perlman 20462 | UC | UC1931262 | Maui, Hawai'i | herb |
| Doryopteris | decipiens | CM Tribble 143 | UC | NA- not yet accessioned | Maui, Hawai'i | silica |
| Doryopteris | decipiens | KR Wood 11555 | UC | UC1930236 | Maui, Hawai'i | herb |
| Doryopteris | decipiens | C Morden 2138 | HAW | HAW04681 | Maui, Hawai'i | herb |
| Doryopteris | decipiens | DD Palmer 1122 | HAW | HAW41408 | O'ahu, Hawai'i | herb |
| Doryopteris | decipiens | DD Palmer 1086 | HAW | HAW41392 | O'ahu, Hawai'i | herb |
| Doryopteris | decipiens | DD Palmer 2863 | HAW | HAW41409 | O'ahu, Hawai'i | herb |
| Doryopteris | decipiens | DD Palmer 1123 | HAW | HAW41394 | O'ahu, Hawai'i | herb |
| Doryopteris | decipiens | CM Tribble 135 | UC | NA- not yet accessioned | O'ahu, Hawai'i | silica |
| Doryopteris | decipiens | DD Palmer 3110 | HAW | HAW41390 | O'ahu, Hawai'i | herb |
| Doryopteris | decipiens | K Mehltreter 1402 | HAW | HAW02420 | O'ahu, Hawai'i | herb |
| Doryopteris | subdecipiens | C Morden 1119 | HAW | HAW04650 | Hawai'i Island, Hawai'i | herb |
| Doryopteris | subdecipiens | KR Wood 12923 | UC | UC1931268 | Kaho'olawe, Hawai'i | herb |
| Doryopteris | subdecipiens | DD Palmer 584 | HAW | HAW41386 | Kaua'i, Hawai'i | herb |
| Doryopteris | subdecipiens | KR Wood 18028 | PTBG | PTBG1000070234 | Kaua'i, Hawai'i | silica |
| Doryopteris | subdecipiens | DD Palmer 597 | HAW | HAW41396 | Kaua'i, Hawai'i | herb |
| Doryopteris | subdecipiens | DD Palmer 763 | HAW | HAW41389 | O'ahu, Hawai'i | herb |
| Doryopteris | subdecipiens | KR Wood 10713 | UC | UC1877540 | O'ahu, Hawai'i | herb |
| Doryopteris | subdecipiens | DD Palmer 764 | HAW | HAW41384 | O'ahu, Hawai'i | herb |
| Doryopteris | subdecipiens | KR Wood 10715 | UC | UC1877541 | O'ahu, Hawai'i | herb |
| Doryopteris | subdecipiens | DD Palmer 3222 | HAW | HAW41388 | O'ahu, Hawai'i | herb |
| Doryopteris | subdecipiens | KR Wood 3169 | UC | UC1871435 | O'ahu, Hawai'i | herb |
| Doryopteris | angelica | CM Tribble 185 | BISH | NA- not yet accessioned | Kaua'i, Hawai'i | silica |
| Doryopteris | angelica | CM Tribble NA | NTBG | NA-living collection | Kaua'i, Hawai'i | silica |
| Doryopteris | angelica | KR Wood 16584 | PTBG | PTBG1000083806 | Kaua'i, Hawai'i | silica |
| Doryopteris | angelica | A Vernon 81 | HAW | HAW39755 | O'ahu, Hawai'i | herb |
| Doryopteris | takeuchii | DD Palmer 1081 | HAW | HAW04485 | O'ahu, Hawai'i | herb |
| Doryopteris | takeuchii | 20200306SCH02 | NA | NA | O'ahu, Hawai'i | silica |
| Doryopteris | collina | J. R. Abbot 16375 | UC | UC1619167 | Santa Cruz, Bolivia | herb |
| Doryopteris | concolor | Meier 4355 | UC | UC1737252 | Carabobo, Venezuela | herb |
| Doryopteris | concolor | Calatayud 4121 | UC | UC2030574 | Cusco, Perú | herb |
| Doryopteris | concolor | Calatayud 2417 | UC | UC1870452 | Cusco, Perú | herb |
| Doryopteris | concolor | EM Zardini 37147 | UC | UC1746417 | Guayaquí, Paraguay | herb |

Continued on next page

Table S1 – continued from previous page

| Genus | Species | Collection Number | Institution | Accession Number | Locality | Material |
|---|---|---|---|---|---|---|
| Doryopteris | concolor | A Bodenghien 2033 | UC | UC1532380 | Lualaba, Democratic Republic of the Congo | herb |
| Doryopteris | concolor | S Perlman 19746 | UC | UC1871439 | Marquesas Islands | herb |
| Doryopteris | concolor | E. Schuettpelz 1396 | MO | MO102497544 | Minas Gerais, Brazil | silica |
| Doryopteris | concolor | Nitta 330 | UC | UC1969724 | Mo'orea, French Polynesia | herb |
| Doryopteris | concolor | Ranker 1939 | UC | UC1784945 | Mo'orea, French Polynesia | herb |
| Doryopteris | concolor | J Game 94/001 | UC | UC1952102 | Rarotonga, Cook Islands | herb |
| Doryopteris | concolor | E. Schuettpelz 1455 | MO | MO102497674 | Rio de Janeiro, Brazil | silica |
| Doryopteris | concolor | Jansen-Jacobs 3561 | UC | UC1612350 | Rupununi, Guyana | herb |
| Doryopteris | concolor | JRI Wood 12276 | UC | UC1788812 | Santa Cruz, Bolivia | herb |
| Doryopteris | concolor | LY Kuo 195 | TAIF | NA-unknown | Taiwan | silica |
| Doryopteris | concolor | KR Wood 10787 | PTBG | PTBG1000023512 | Ua Huka, French Polynesia | silica |
| Doryopteris | lorentzii | J. Prado 2126 | DUKE | NA-unknown | São Paulo, Brazil | silica |
| Doryopteris | nobilis | UCBG 69.0032 | UCBG | UCBG 69.0032 | Brazil | silica |
| Doryopteris | patula | UCBG9 7.0581 | UCBG | UCBG 97.0581 | Brazil | silica |
| Doryopteris | pentagona | E. Barbosa 276 | UC | UC1735287 | Parana, Brazil | herb |
| Doryopteris | cf. varians | Mynssen CM 873 | UC | UC1877755 | Espirito Santo, Brazil | herb |
| Doryopteris | varians | E. Schuettpelz 1454 | UC | UC1950769 | Rio de Janeiro, Brazil | silica |
| Lytoneuron | crenulans | J. Prado 2135 | UC | UC1950748 | Santa Catarina, Brazil | silica |
| Lytoneuron | lomariacea | E. Schuettpelz 1462 | MO | 102497564 | São Paulo, Brazil | silica |
| Lytoneuron | paradoxa | E. Schuettpelz 1442 | MO | 102497556 | Rio de Janeiro, Brazil | silica |
| Lytoneuron | quinquelobatum | P. Fischi 3898 | SPF | NA-unknown | Rio de Janeiro, Brazil | silica |
| Ormopteris | gleichenioides | E. Schuettpelz 1374 | MO | 102497532 | Minas Gerais, Brazil | silica |
| Ormopteris | pinnata | E. Schuettpelz 1377 | MO | 102497538 | Minas Gerais, Brazil | silica |
| Ormopteris | riedelii | E. Schuettpelz 1387 | MO | 102497604 | Minas Gerais, Brazil | silica |

### S2 matK phylogeny

We additionally sequenced matK amplicons from a subset of individuals using PacBio's Sequel II sequencing platform. We processed the sequences using the PURC pipeline (Schafran et al., 2023; Rothfels et al., 2017) and inferred a maximum likelihood phylogeny with `IQ-TREE multicore version 1.6.12` under default settings, except with 10,000 ultrafast bootstrap replicates and inferring the best-fitting model with `ModelFinder` (Nguyen et al., 2015; Kalyaanamoorthy et al., 2017). The resulting phylogeny was largely concordant with the results from our nuclear genome-wide analysis presented in the main text, though with poorer resolution: Hawaiian *Doryopteris* is nested within *D. concolor* and sister to other *Doryopteris* spp. *Doryopteris* is monophyletic and sister to a clade of *Lytoneuron* and *Ormopteris*, which are reciprocally monophyletic. Hawaiian individuals show very little resolution, but do not appear to form clades based on species identity.

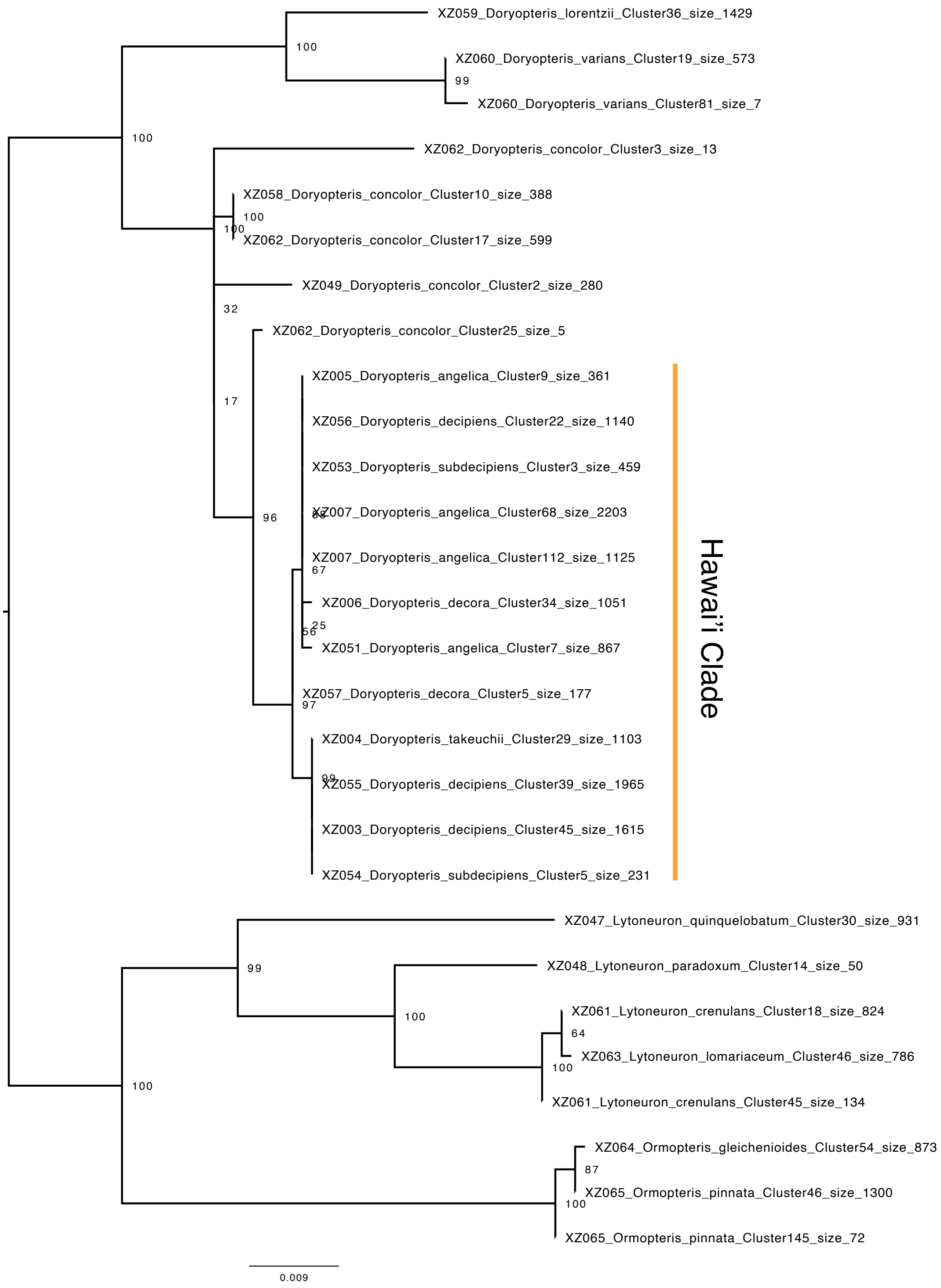


**Figure S1:** A phylogeny of *Doryopteris* and outgroups using a single matK chloroplast locus. Numbers at nodes correspond to ultrafast bootstrap support values.

#### S3 ASTRAL2 full tree

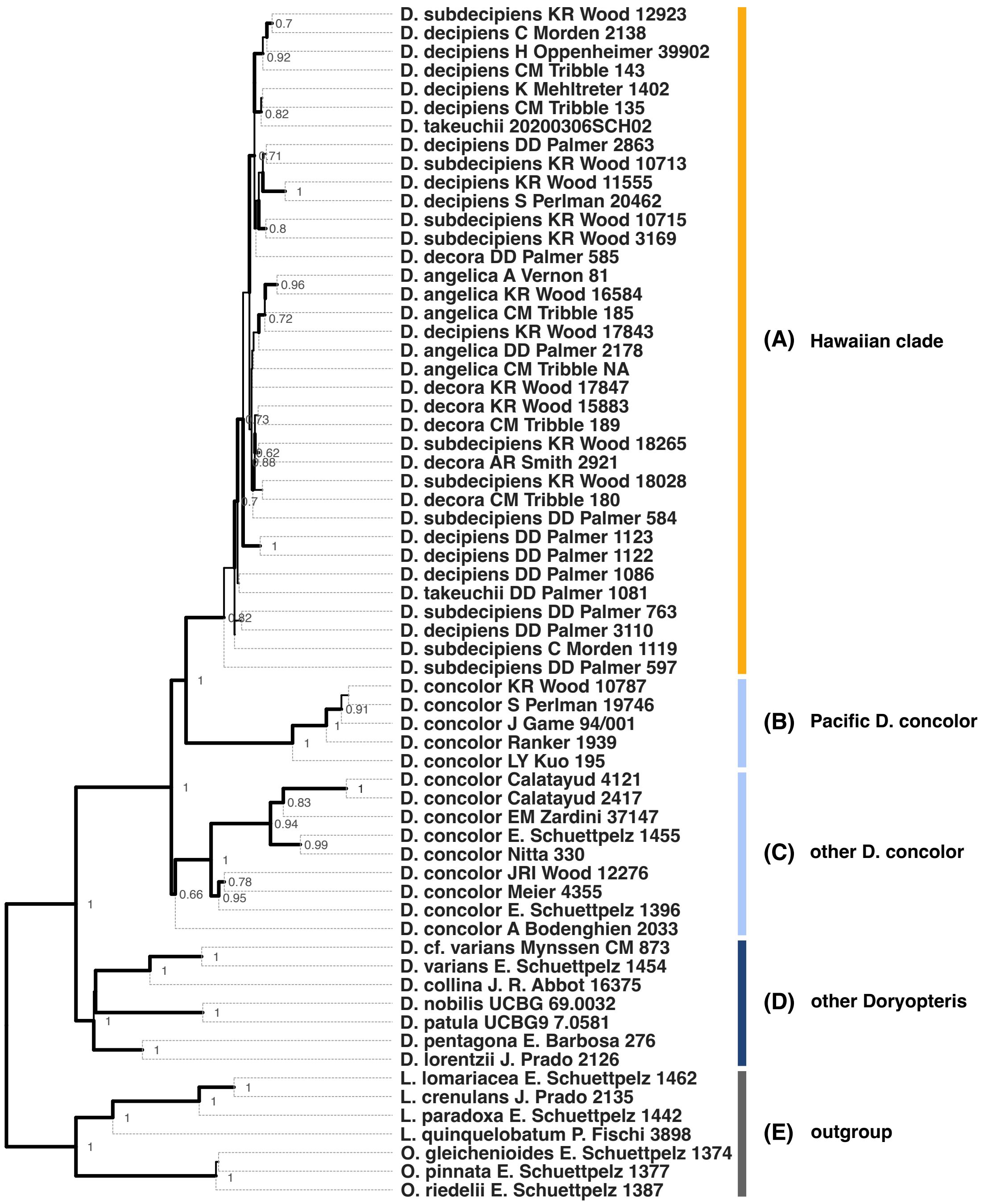


**Figure S2:** Full tree of Fig 2 in main text. A phylogeny of *Doryopteris* and outgroups using the GoFlag probe set inferred using ASTRAL-II. Numbers at nodes correspond to posterior support values. Only support ¿0.6 is shown.
